# Time-resolved proteomic analysis of hepatitis B virus infection

**DOI:** 10.64898/2026.02.24.707876

**Authors:** Mohammed A. Akbor, Medhi Boutasbih, Alexander Koenig, Eun-Ji Jo, Iolanda Vendrell, Svenja S. Hester, Senko Tsukuda, Monica M. Olcina, Marc P. Windisch, Jane A. McKeating, Benedikt M. Kessler, Andreas Damianou

## Abstract

Hepatitis B virus (HBV) infection is a major cause of chronic liver disease and hepatocellular carcinoma, yet its impact on host proteome dynamics during early infection remains poorly defined. Here, we performed time-resolved label-free quantitative (LFQ) proteomics to characterise the host response to HBV infection in human HepG2 hepatoma cells. Across five time points spanning the first three weeks of infection, we quantified 5,365 unique proteins, including two HBV proteins (C/preC-derived and S), revealing subtle yet coordinated changes in the hepatocellular proteome. Early infection was characterised by selective alterations in cytoskeletal organisation, extracellular matrix components, and focal adhesion pathways, alongside enrichment of complement-associated proteins. Notably, we identified a transient, specific increase in the abundance of complement component C5 without a corresponding change in C3, indicating differential regulation of complement pathway members during infection. These findings define a temporally resolved proteomic landscape of HBV–host interactions and uncover previously underappreciated complement dynamics consistent with a selective and restrained host response. Together, this provides a framework for mechanistic studies of early viral infection and host adaptation.

## INTRODUCTION

Hepatitis B virus (HBV) is a prototypic member of the *Hepadnaviridae* family that is frequently associated with a persistent infection of the liver. Chronic hepatitis B (CHB) is a global health problem, with approximately 2 billion people exposed to the virus during their lifetime, resulting in ∼250 million chronic infections. CHB is associated with an increased risk of progressive liver diseases, including cirrhosis and hepatocellular carcinoma, contributing to an estimated 820,000 deaths annually^1^. HBV infects hepatocytes through the sodium taurocholate co-transporting polypeptide (NTCP) receptor ^2,3^ and persists in the nucleus as a covalently closed circular DNA (cccDNA) complexed with histones. This cccDNA is transcribed by host RNA Polymerase II to yield six major overlapping transcripts: pre-core (pC) encoding E antigen; pre-genomic (pgRNA) encoding core protein (C) and polymerase; the preS1, preS2 and S RNAs encoding the surface glycoproteins (S); and the X transcript encoding regulatory X protein.^4^. Our recent long-read RNA sequencing studies show that transcripts encoding S dominate the viral transcriptome, while genomic-length pC and pgRNA represent a relatively minor component (∼8%)^5,6^. As the pC and C proteins are translated from the same core open reading frame (ORF), they share an amino acid sequence. The pC protein is translocated into the endoplasmic reticulum and proteolytically processed to generate the secreted hepatitis B e antigen (E), whereas C protein remains intracellular and assembles into viral nucleocapsids^7^.

Current nucleoside analogue therapies rarely cure infection as they have limited impact on the long-lived HBV cccDNA reservoir^8^. Several new therapeutic strategies, including HBV-specific small interfering RNAs (siRNAs) and epigenetic modifying agents, are in development to cure infection^9-11^. While HBV replication has been extensively studied at the molecular level, the global impact of infection on the host cellular proteome remains partially understood. As viral replication is fundamentally intertwined with host cellular pathways, defining viral-driven changes in the host proteome can identify mechanisms of viral persistence and potential new therapeutic targets.

Viruses are obligate intracellular pathogens that depend on the host cell to complete their infectious life cycle. Throughout evolution, they have adapted strategies to interact with and manipulate host cell processes to support their replication and persistence. Although several transcriptomic studies have provided important insights into HBV host interplay using immortalised cell lines and primary human hepatocytes^12-14^, RNA abundance may not always associate with protein levels or activity, underscoring the need for proteome-wide analysis. Earlier proteomic reports studied the HBV producer cell line HepG2.2.15 or hepatoma cells transfected with infectious HBV DNA plasmids to evaluate the host response to viral infection^15-18^. Although these studies identified differentially abundant proteins (DAPs) involved in lipid metabolism, proliferation, cytoskeletal, and mitochondrial functions, they reported only a limited number of DAPs. Advanced mass spectrometry-based proteomic techniques provide more powerful, quantitative tools for studying proteome-wide alterations. Yuan et al. reported extensive proteome-wide changes, identifying >1,300 DAPs in HepG2.2.15 cells compared with parental HepG2 cells, and noted modulation of NADH dehydrogenase, ATP synthase, and cholesterol metabolism during viral infection. Recent studies sought to investigate host responses *in vivo* in a fibrotic context, utilising both HBV-transgenic mouse models and clinical samples from patients diagnosed with CHB, providing new insights into virus-driven changes in metabolic pathways and oxidative stress effectors^19,20^. It’s noteworthy that published *in vitro* models have major limitations: the absence of an authentic infection model and the use of HepG2.2.15 cells, which were generated over 30 years ago and have been maintained in continuous culture, making parental HepG2 cells an imperfect comparator^21^. As such, the early events defining the virus-host interplay remain poorly characterised. Since the discovery of NTCP as the main entry receptor for HBV, NTCP-expressing HepG2 cell lines have emerged as a more biologically relevant model that supports the full replicative life cycle, from entry to egress.

We previously developed and validated the HepG2-NTCP-sec+ cell line, a replication- and infection-competent hepatocyte model that supports the complete HBV life cycle from viral entry through virion production and release^22^. Using this well-characterised infection model^22^, we investigated early events following HBV infection through a time-resolved quantitative mass spectrometry approach to characterise proteome-wide alterations in hepatocellular biology. We identified dynamic alterations in cytoskeletal organisation, extracellular matrix (ECM) components, and complement-associated proteins in response to HBV infection. Together, these findings establish a time-resolved, hypothesis-generating proteomic atlas of HBV–host interactions during the first three weeks of infection, providing a framework to guide future mechanistic studies.

## METHODS

### Cell culture and HBV infection

HepG2-NTCP-sec+ cells were cultured as described previously (König et al., 2019). Briefly, 5 × 10^6 cells were seeded in T-75 flasks in culture medium containing DMEM (Welgene LM001-07), 2.5% DMSO (Sigma D2650), 10% FBS (Gibco 16000-044), 1% L-glutamine (Gibco 25030-081) and penicillin/streptomycin (Gibco 15140-122). Three independent flasks were used as biological replicates for each experimental condition. Cells were maintained in DMSO-containing medium from the time of seeding, and HBV or mock inoculation was initiated approximately 24 h after plating. Cells were inoculated for 16 h with HepAD38-cell-harvested HBV genotype D (2,500 genome equivalents [GEq]/cell, 5 × 10^8 GEq/mL) in culture medium supplemented with 4% PEG8000; mock-infected cultures received culture medium containing 4% PEG8000 without HBV. Each biological replicate was independently inoculated.

The end of the 16-h inoculation period was designated as day 1 post-infection (1 dpi) and cells were washed three times with prewarmed PBS. Cultures were assigned to subsequent time points and were maintained until 4, 7, 11, or 21 dpi, with medium changes every 3–4 days. At each collection time point, cells were washed three times with prewarmed PBS, and monolayers collected in 2 mL prewarmed PBS into 50-mL conical tubes and diluted to 40 mL with prewarmed PBS. Cells were centrifuged at 1,500 × g for 5 min, the supernatant discarded and pellets resuspended in 1 mL prewarmed PBS in 1.5-mL tubes. Samples were centrifuged at 3,000 × g for 5 min and the resulting cell pellets frozen in liquid nitrogen.

### Immunofluorescence of HBsAg ang HBcAg

Immunofluorescence staining was performed as previously described^22^. Cells were washed once with PBS and fixed with 4% paraformaldehyde (Biosesang, Seongnam-si, South Korea). Fixed cells were washed four times with PBS and permeabilized with 0.075% Triton X-100 in PBS for 20 min and washed with PBS three times. Cells were incubated for 16 h at 4ºC with blocking buffer (5% bovine serum albumin in PBS, Sigma-Aldrich, St Louis, USA). Cells were incubated with rabbit anti-HBcAg antibody (C1-5 mouse monoclonal, Santa Cruz, USA) or anti-HBsAg antibody diluted in 5% bovine serum albumin in PBS for 1 hour, repeatedly washed, and incubated with secondary antibodies conjugated with Alexa Fluor 488 (Thermo Fischer, Waltham, USA). Nuclei were stained with Hoechst33342 (Thermo Fischer, Waltham, USA).

### HBV DNA quantification

DNA from HBV infected and mock-infected HepG2-NTCPsec+ were extracted using the DNeasy Blood & Tissue kit (Qiagen, Hilden, Germany) and analyzed for HBV cccDNA as previously described^23^. Briefly, primers 5⍰-CGTCTGTGCCTTCTCATCTGC-3⍰, 5’-GCACAGCTTGGAGGCTTGAA-3’, and a fluorogenic 5’-end nuclease probe 5’-FAM-CTGTAGGCATAAATTGGT-(MGB)-3’ were used. TaqMan Universal PCR master mix (Thermo Scientific, Waltham, USA) with 0.2 μM primer, 0.1 μM probe, and 2 μL extracted DNA were prepared and analyzed with a VIIA7 real-time PCR detection system (Thermo Scientific, Waltham, USA).

### Immunoblotting

Immunoblotting was performed as previously described^22^. Briefly, cells (1 × 10^^6^) were washed with ice-cold PBS and lysed in RIPA buffer supplemented with protease and phosphatase inhibitors. A total of 50 µg protein was resolved by SDS–PAGE on acrylamide gels and transferred to nitrocellulose membranes. Membranes were blocked in 5% milk and incubated with primary antibodies against HBc and GAPDH, followed by the appropriate conjugated secondary antibodies.

### Mass spectrometry sample preparation

Buffers for resuspending peptides were prepared using HPLC-grade reagents. Cell pellets were resuspended in RIPA lysis and extraction buffer supplemented with 0.1μL of Benzonase® Nuclease. Lysates were sonicated in a water bath for 5 mins. SDS was added to the lysates to a final concentration of 5%. Samples were then processed using the S-Trap™ micro/midi spin column digestion protocol (C02-micro/C02-midi, Protifi LLC), depending on protein amounts. Manufacturer instructions were followed, using 5 mM tris(2-carboxyethyl)phosphine (TCEP) and 20 mM iodoacetamide (IAA) as the reducing and alkylating agents, respectively. Proteins were digested overnight with TPCK-treated trypsin (20233, ThermoFisher) at the ratio recommended by S-Trap instructions (1:25, protein:trypsin). Resultant peptides were eluted and dried using a SpeedVac concentrator centrifuge. Dried peptides were resuspended in an appropriate buffer for mass spectrometry (MS) analysis or peptide-level enrichment.

### Mass spectrometry analysis

Peptides were resuspended in 5% formic acid and 5% DMSO and analysed by liquid chromatography using an Ultimate 3000 UHPLC coupled to a Q-Exactive (QE) mass spectrometer (ThermoFisher). Peptides were trapped on an Acclaim™ PepMap™ 100 C18 HPLC Column (100µm x 2mm, 5µm particle size, 164750, ThermoFisher) and separated on an EasySpray PepMap RSLC column (75µm i.d. x 2µm x 50cm, 100 Å, ThermoFisher) using a linear gradient of 5% to 35% solvent B (0.1% FA in ACN and 5% DMSO), with a length of 120 min and a flow rate of 250nL/min. The raw data were acquired in the mass spectrometer in data-independent acquisition (DIA) mode. Full scan MS spectra were acquired in the Orbitrap (inclusion list with scan range 495-995 m/z, 20 m/z increments, with an overlap of +/-2 Daltons, resolution 35000, AGC target 3e6, maximum injection time 55ms). After the MS scans, peaks were selected for higher-energy collisional dissociation (HCD) fragmentation at 28% of normalised collision energy (NCE) / stepped NCE. HCD spectra were also acquired in the Orbitrap (resolution 17500, AGC target 1e6, isolation window 20 m/z). Samples were analysed sequentially with replicate 1 run before replicate 2 and subsequently replicate 3. Virus infected samples were analysed in chronological order (2, 4, 7, 1 and 21 dpi) followed by the corresponding mock samples, with a blank injection between each sample. HeLa peptide samples were analysed to monitor mass spectrometer performance throughout the acquisition.

### Bioinformatics analysis

Raw DIA MS data from proteomic analysis were analysed using DIA-NN v1.9.2^24^ with library-free search and default settings (trypsin/P protease, 1 missed cleavage, FASTA digest for library-free search, N-term M excision as a variable modification, and C carbamidomethylation as a fixed modification). Match between runs (MBR) was enabled and protein inference was performed on the species-specific gene level. Data were searched against a reference proteome that included human proteins (Proteome ID: UP000005640) and HBV genotype D proteins (Proteome ID: UP000007930), downloaded from UniProt on 1st October 2023. Searches were performed with and without common Repository of Adventitious Proteins (cRAP), since several cRAPs have been previously detected in hepatocytes and/or liver tissue^25^. The data analysis focused on the precursor and unique gene matrices generated by DIA-NN (FDR filter was 0.01). Bioinformatic analysis and visualisation of data were primarily performed in RStudio^26^ 2023.03.0+386 (R, v4.3.2) using ggplot2^27^. MS raw data have been uploaded to the ProteomeXchange Consortium via the PRIDE partner repository^28^ with the dataset identifier PXD074747.

### Quality control

Data quality control (QC) involved principal component analysis (PCA), calculation of Pearson correlation coefficients (PCCs), and inspection of the distribution of intensity values. Precursors corresponding to the same peptide sequence were collapsed by summing intensity values. Proteins/peptides were filtered by removing those not identified in 3 biological replicates at any time point or infection treatment. Data were log2-transformed and normalised by median subtraction within each replicate. The remaining missing values were imputed from a down-shifted normal distribution (width = 0.3, shift = -1.8) to emulate low-abundance proteins.

### Differential abundance analysis

For differential abundance analysis, the moderated t-test was used to shrink sample variances and increase statistical power using an empirical Bayes approach. The Benjamini-Hochberg procedure (BH) for false discovery rate (FDR) control was applied to the moderated t-test p-values. HBV-infected samples were compared to mock-infected samples for each time point, and results were visualised using volcano plots.

### Weighted correlation network analysis

WGCNA was performed in R using the corresponding R package^29^. Log2-transformed, median-normalised, and imputed data were scaled, resulting in ‘Z-scored’ values to use as the WGCNA input. Soft-thresholding optimisation **(Fig. S7)** was performed to determine an optimal power for a signed network. Parameters included TOMType = ‘signed’, networkType = ‘signed’, corType = ‘bicor’, power = 12, minModuleSize = 20, reassignThreshold = 0, mergeCutHeight = 0.2, and maxPOutliers = 0.05. Module robustness was measured using the ‘modulePreservation’ function using the dataset randomly divided into training and testing datasets, using the same parameters for network construction with 1000 permutations. Fréchet distances between eigen proteins corresponding to HBV-infected and mock-infected samples were measured for resultant modules from WGCNA. Module eigen values were mean averaged for each time point and treatment condition for visualisation purposes

### Functional enrichment

Overrepresentation analysis (ORA) used the enrichGO and enrichKEGG functions from the clusterProfiler package^30^, using genes corresponding to proteins identified in the datasets as the background, rather than the whole genome. Gene set enrichment analysis (GSEA) was performed using the clusterProfiler GSEA function, with Log2FC as the ranking metric.

STRING network analysis was performed in Cytoscape^31^, using the RCy3 package^32^ in R and the Cytoscape StringApp plugin^33^.

### Multiple sequence alignment

Proteins of interest were queried against the UniProt database to identify corresponding homologs in Bos taurus (taxon ID: 9913). Human and bovine proteins that shared the most similar lengths were aligned using the ClustalOmega algorithm. Detected peptides were then mapped onto and compared to bovine versions in the sequence alignment.

## RESULTS

In this study, we investigated the dynamic alterations in the proteome of HepG2-NTCP-sec+ cells during HBV infection^22^. As a control, parallel cultures were mock-infected to ensure that any observed changes could be attributed to the viral infection rather than experimental conditions. Infected cells were sampled at five time points: 1, 4, 7, 11, and 21 days post-infection (dpi), in triplicate to ensure robust data acquisition and reproducibility **(Fig. 1A)**. HBV infection was assessed by PCR measurement of intracellular HBV DNA, immunoblotting of HBc, and immunofluorescent detection of HBc and HBsAg **(Fig. S1)**. Using high-resolution mass spectrometry, we identified and quantified 5,365 unique proteins across all experimental samples. We observed a temporal decline in the number of proteins identified in both mock and HBV-infected samples, suggesting that the culture conditions per se induced progressive alterations in the cellular proteome **(Fig. 1B)**. Importantly, the consistent detection of viral proteins in all infected samples **(Fig. S1A, Fig. S2A)** confirmed time-dependent HBV infection.

**Figure 1:**
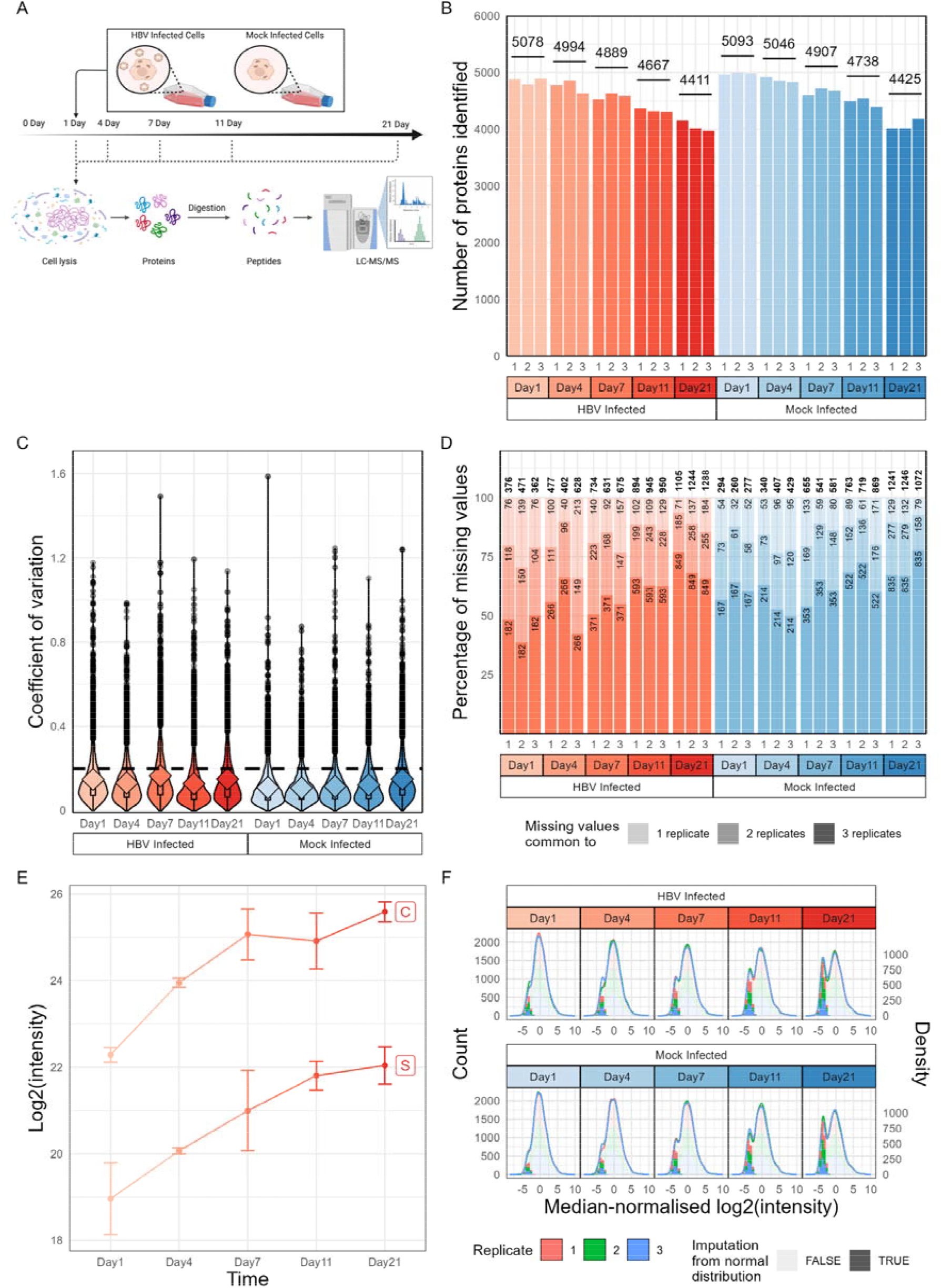
Quality control assessment of proteomic analysis of mock- and HBV-infected HepG2 cells. (A) Schematic representation of the proteomic study, detailing the infection of HepG2-NTCP-sec+ cells, followed by sample collection at multiple time points, protein digestion, and mass spectrometry analysis. (B) Bar chart showing the number of unique proteins identified in each sample. (C) Coefficient of variation (CV) analysis per biological condition, illustrating the experiment’s reproducibility. (D) Distribution of missing proteins per sample, highlighting the consistency across replicates. (E) Log_2_(intensity) of HBV Core (C) and Surface (S) proteins. HBV proteins were not detected in the mock-infected samples. (F) Log_2_-transformed, median normalized distributions of protein intensities for each time point and condition.

To assess the reproducibility and technical variation of the proteomic data, we calculated the coefficient of variation (CV) for each experimental condition. The low CV values across the triplicate samples highlight data reproducibility **(Fig. 1C)**. In addition, we evaluated the consistency of missing protein identifications across replicate samples; more than half of the missing values were absent in all three replicates **(Fig. 1D, F)**. Screening for HBV-encoded proteins identified two viral proteins, C and S, present only in infected samples, whose accumulation over time reflects ongoing replication and viral amplification within infected cells **(Fig. 1E, Fig. S2A)**. The detected viral C peptides mapped to the shared preC/C coding sequence, from which both C and the pC-derived E antigens are translated **(Fig. S2B)**. As the identified tryptic peptides are common to both gene products, our experimental mass spectrometry workflow cannot distinguish C from preC-derived proteins. Accordingly, we refer throughout this manuscript to the detected pC/C viral peptides collectively as the C protein. Subsequently, we applied stringent filtering criteria to include proteins identified in all three replicates within at least one biological condition **(Fig. 2A)**. Missing values were imputed from a downshifted normal distribution **(Fig. 2A)**, enabling us to proceed with downstream comparative analyses. As expected, the requirement for imputed values increased at later time points, in line with the decreasing number of proteins over time **(Fig. 1F)**.

**Figure 2:**
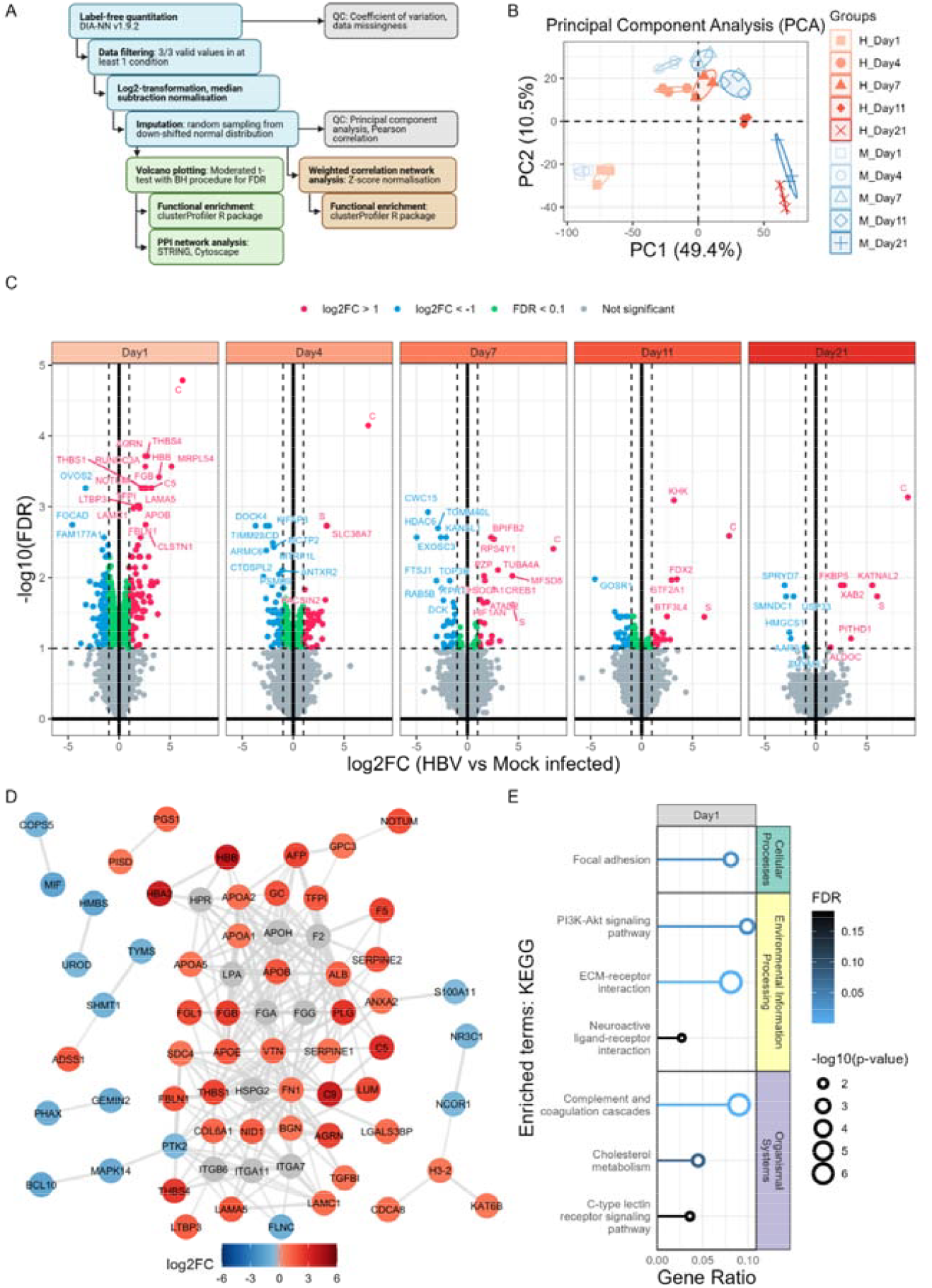
Bioinformatic analysis of mock- and HBV-infected HepG2 cells at early time points reveals alterations in complement and coagulation cascade and extracellular matrix interactions. (A)Schematic representation of the bioinformatic workflow following mass spectrometry analysis. (B)Principal component analysis of log_2_-transformed and median-adjusted data containing imputed values. (C) Volcano plots showing differentially abundant proteins in HBV-infected cells at each time point. Upregulated proteins (FDR < 0.1, log_2_FC > 1) and downregulated proteins (FDR < 0.1, log_2_FC < - 1) are shown in red and blue, respectively. Green points represent proteins with significant FDR but small fold changes (FDR < 0.1, abs(log_2_FC) < 1). (D) Protein–protein interaction network of significantly upregulated and downregulated proteins at 1 dpi. Node colour represents the log_2_FC in protein abundance relative to mock controls, as derived from quantitative proteomics. Grey nodes represent additional interactors (not identified as DAPs), as determined by STRING. Edges denote high-confidence functional and/or physical associations (minimum required interaction score > 0.7). Isolated (unconnected) nodes were removed for clarity. Interaction data were retrieved from the STRING database (v12.0) using the stringApp in Cytoscape (v3.10.0). (E) Functional enrichment analysis of upregulated and downregulated proteins at 1 dpi. Enriched terms for the KEGG pathways dataset are shown, using the cellular proteome as background.

Principal component analysis of the log_2_-transformed, median-normalised, and imputed data showed separation of experimental conditions by HBV infection and dpi **(Fig. 2B)**, suggesting that both infection and prolonged culturing influenced the cellular proteome. The progressive shift along principal component (PC) 1 reflected a time-dependent remodelling of the proteome, whereas separation along PC 2 distinguished HBV infection from mock conditions. This dual effect suggests that infection imprints a distinct proteomic trajectory on the culture-driven background changes.

Volcano plots comparing HBV- and mock-infected cells at each time point revealed groups of DAPs **(Fig. 2C)**. The greatest number of DAPs was observed at 1 dpi, suggesting that the initial stages of HBV infection result in strong proteomic perturbations **(Fig. S3A)**. To further characterise the DAPs at each time point, we performed STRING-based protein–protein interaction (PPI) analysis. At 1 dpi, when the largest number of DAPs was detected, the network displayed the highest degree of connectivity **(Fig. 2D)**, compared to 4 and 7 dpi **(Fig. S5)**. The 1 dpi network comprised a densely interconnected cluster of complement proteins, including C5 and C9, coagulation-related proteins such as fibrinogen subunits (FGA, FGB, FGG) and plasminogen activator inhibitor 1 and 2 (SERPINE1 and SERPINE2), lipid binding proteins such as the apolipoprotein (APO) family of proteins, and extracellular matrix proteins, including Collagen alpha-1(VI) chain (COL6A1), fibulin-1 (FBLN1), and fibronectin (FN1). These interactions demonstrate that the earliest response to HBV infection involved a coordinated modulation of immune and structural protein categories. Down-regulated proteins were fewer in number, were less strongly connected, and included metabolic enzymes and regulatory proteins such as B-cell lymphoma/leukaemia 10 (BCL10), COP9 signalosome complex subunit 5 (COPS5), and focal adhesion kinase 1 (PTK2), all of which were positioned at the periphery of the network. The high density of nodes and edges highlights the scale of infection-driven host remodelling at early time points following infection.

KEGG pathway enrichment analysis of the DAPs at 1 dpi revealed four major pathways: complement and coagulation cascades, ECM–receptor interactions, proteoglycans in cancer, and focal adhesion **(Fig. 2E)**. Complement and coagulation cascades showed the strongest enrichment signal, consistent with the dense STRING network cluster of C5, C9, fibrinogen subunits, and related factors. ECM– receptor interaction and focal adhesion pathways were also enriched, driven by collagens, integrins, fibronectin, and other structural proteins. These results confirm that the earliest stage of infection is characterised by coordinated alterations in immune and ECM pathways, consistent with the STRING connectivity patterns. In contrast, there were modest changes in interferon-stimulated gene (ISG) expression, with 8 proteins showing differential abundance at specific time points **(Fig. S4)**.

In addition to the time-dependent patterns, we examined proteins altered throughout the entire infection time course **(Fig. S3B)**. Five proteins were identified as differentially expressed between 1 and 11 dpi: the viral C and S proteins, bactericidal/permeability-increasing fold-containing family B member 2 (BPIFB2), FN1, and pregnancy zone protein (PZP). This highlights the transient nature of the proteomic changes and a limited set of infection-associated markers.

To complement the functional enrichment analysis, gene set enrichment analysis (GSEA) was performed on a ranked list of genes based on corresponding protein log_2_(fold change) (log_2_FC; HBV versus mock infection) at each time point **(Fig. 3A)**. Identified gene sets demonstrated higher significance (lower FDR) at 1 dpi. Nevertheless, the hallmark gene sets ‘epithelial mesenchymal transition’ (EMT) and ‘coagulation’ (and similar GOBP gene sets) showed decreasing normalised enrichment scores (NES) over time, while ‘DNA repair’ increased. GOMF gene sets corresponding to glycosaminoglycan- and collagen-binding, as well as extracellular matrix, also showed elevated NES at 1dpi, with NES decreasing as infection progressed. GSEA plots depicting the running enrichment scores, leading edges, and distributions of ranking metrics for notable pathways confirm differential enrichment of coagulation, DNA repair, and EMT pathways across the infection time course **(Fig. 3B, C)**. KEGG ECM receptor interactions demonstrated a similar enrichment trend, reinforcing the notion of early modulation of ECM-related processes at the onset of HBV infection. In summary, these analyses show that pathways altered at early time points become less pronounced as infection progresses.

**Figure 3:**
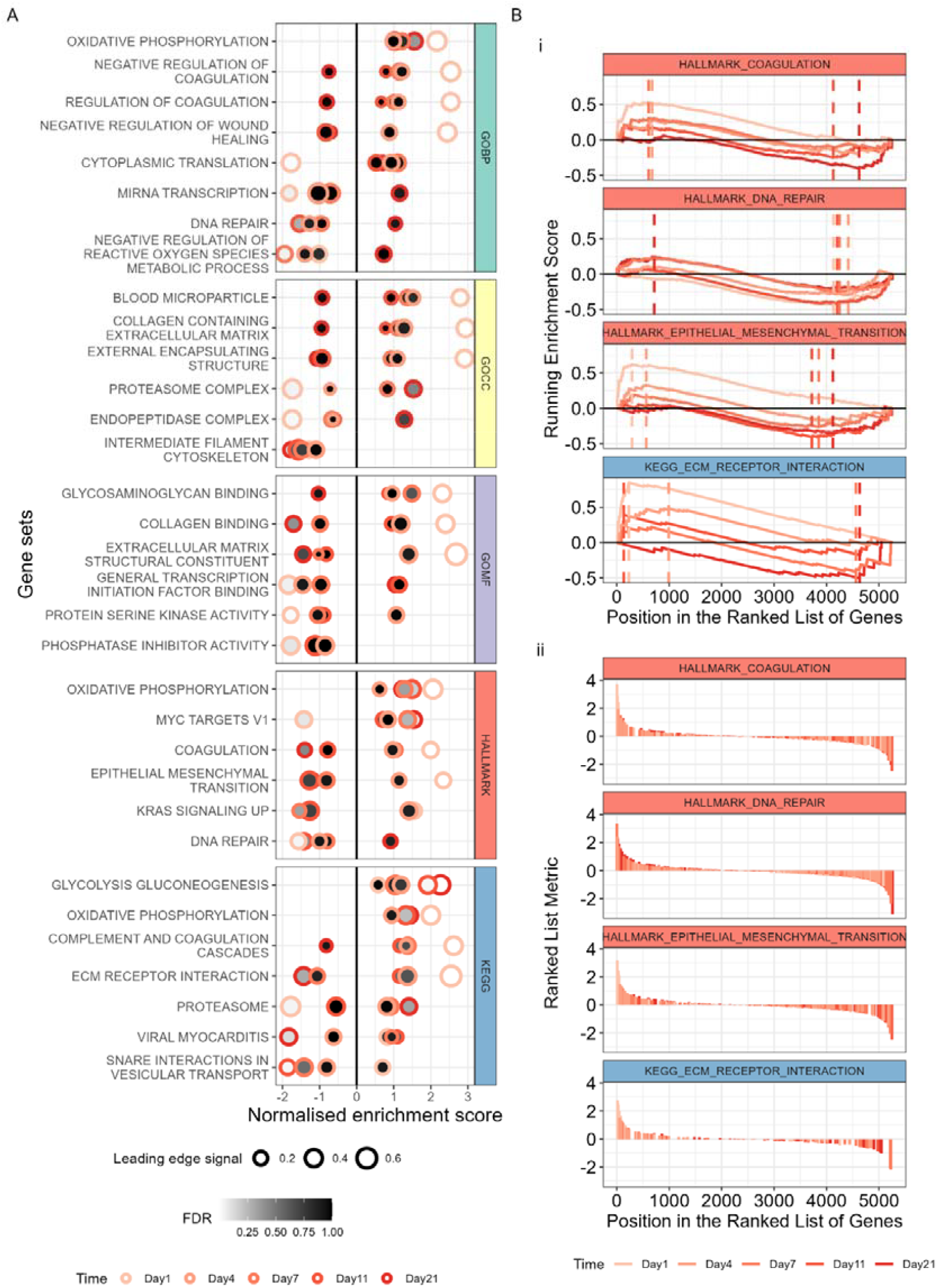
GSEA of the HBV infection time course proteomics identifies hallmark, GO, and KEGG gene sets. (A) Genes corresponding to proteins identified in proteomics data were ranked based on log_2_FC. (B) GSEA plots showing running enrichment score (i) and ranked list metric (ii) on the y-axis. Complement- and coagulation-related processes were consistently detected in the functional enrichment analyses. Notably, we observed a transient elevation of complement C5 protein during early HBV infection **(Fig. 4A)**. Sequence comparison with the corresponding bovine homologues showed that most peptides contributing to the complement- and extracellular-associated proteins were unique to human protein sequences **(Fig. S6A-B)**. This was selective for certain components of the complement pathway, as we did not observe a similar statistically significant increase in C3 abundance **(Fig. S6D-E)**. Consistent with previous studies^34^, we observed differences in the peptide distribution across C5 at 4 dpi and subsequent time points **(Fig. 4A)**. The annotated processing sites shown in Fig. 4B provide structural context for the observed peptide distribution. However, our bottom-up proteomic workflow does not directly distinguish cleaved from intact C5; therefore, these data cannot establish C5 proteolytic processing.

Weighted correlation network analysis (WGCNA) was performed on z-scored data, which standardised protein abundances, to identify protein modules with similar temporal profiles **(Fig. S7)**. Module preservation was measured by randomly splitting the data into training and testing datasets for permutation analysis. Only module 5 demonstrated a lower median rank than the ‘gold’ module (representative random module) and a Z-summary statistic less than 2, suggesting this module is unlikely to be preserved **(Fig. S7)**. The other modules demonstrated median ranks higher than the ‘gold’ module, and moderate-to-high module conservation (Z-summary values between 4.5 and 54), indicating greater conservation and biologically-relevant modules. The majority of proteins exhibited similar abundance profiles in the mock- and HBV-infected samples and clustered accordingly **(Fig. 5A)**. However, some modules showed infection-dependent protein abundance profiles, as indicated by higher Fréchet distances between module eigen protein profiles corresponding to mock and HBV-infected samples **(Fig. 5B)**. Proteins in modules 4 and 6 demonstrated unique abundance profiles that indicate reduced protein abundance during HBV infection, whereas module 5 demonstrated lower protein abundance at 1 dpi and higher after 4 dpi upon HBV infection compared to mock infection **(Fig. 5B)**. Functional enrichment analysis linked these proteins to secretory organelles, including the endoplasmic reticulum and Golgi apparatus, as well as mitochondrial components **(Fig. 5C)**. Gene sets associated with glycosylation were also enriched in module 4, which demonstrated lower trajectories in HBV-infected samples, suggesting reduced abundances for proteins associated with protein modifications typically attached to secreted proteins. Despite lower module preservation, module 5 demonstrated enrichment for cytokine production, consisting of the proteins BCL10 and MALT1, two key mediators of NF-KB signalling pathway.

## DISCUSSION

Our study shows that HBV infection elicits a transient remodelling of the host proteome within the first day of infection, dominated by complement-associated changes and extracellular matrix remodelling. In parallel, the reduced abundance of secretory pathway proteins suggests that infection may alter ER–Golgi-associated processes, potentially influencing viral replication and host immune responses. However, the biological significance and functional consequences of these proteomic changes are beyond the scope of the current study and would require further investigation. ECM alterations could be associated with viral infection of HepG2 cells and may contribute to tissue-remodelling processes associated with chronic HBV infection, including fibrosis^3,35,36^; therefore, functional enrichment analyses highlighting GAG- and ECM-binding proteins may support immune evasion strategies that are worth investigating. These findings provide a temporal framework for understanding how HBV reshapes the host proteome during early stages of infection. Notably, these coordinated host responses were most pronounced at 1 dpi and may reflect early events associated with HBV attachment, entry and NTCP-dependent signalling. Viruses commonly interact with host proteins during infection, some of which can be incorporated into viral envelopes or capsids^37^. Such early virus–host interactions may contribute to the more pronounced proteomic changes observed at the earliest times post-infection. Further studies using antiviral agents with well-characterised mechanisms of action are warranted to delineate the specific stages of the viral life cycle responsible for driving these changes in the host proteome.

This temporal pattern suggests that HBV induces an early but transient change in the host proteome, consistent with its well-described ‘stealth’ behaviour in hepatocytes. Previous proteomic studies used a variety of experimental models, including HBV-infected HepG2-NTCP^38-40^; Huh-7 cells^41,42^; HBV-transfected cells^43^; and HBx transgenic mice^44^. Each of the models reflects a different aspect of HBV infection biology, with HepG2 cells, originally derived from a hepatocellular carcinoma, potentially representing an early hepatocyte differentiation state. Major hallmarks uncovered in our study include extracellular matrix remodelling, cytoskeletal rearrangements and focal adhesion pathways in HBV-infected samples, consistent with earlier studies^45,46^. We noted transient elevation and modulation of complement components, such as C5, within the first few days of infection. This observation is notable given that complement-related protein enrichment, which predominantly occurs in the extracellular compartment, has been implicated in modulating HBV-associated liver inflammation and coagulation responses *in vivo*^37^. In a murine model of virus-induced fulminant hepatitis, C5a receptor (C5aR) signalling was shown to contribute to inflammatory and pro-coagulant responses that are associated with disease severity^47^. Non-immune cell-based expression of complement C3a and C5a has been described in the context of liver injury and regeneration^48^. IL-6 expression by hepatocytes exposed to HBV can lead to complement C3 production via hepsin-HBx interactions^49^. C5a was described as a fibrosis and cirrhosis marker in chronic hepatitis B^50^. Our peptide mapping data are consistent with, but do not directly demonstrate, selective C5 processing in the context of HBV infection **(Fig. 4)**. Importantly, sequence comparison with the corresponding bovine homologues indicated that the majority of peptides derived from complement- and extracellular-associated proteins were human-specific **(Fig. S6)**, reducing the likelihood that these signals originated from FBS-derived proteins. The biological relevance of this observation warrants further investigation and may reflect an aspect of the host innate antiviral defence mechanism. Given the central role of type I and III interferons (IFNs) in host antiviral defence, mediated principally through the induction of ISGs, it remains uncertain whether a robust ISG programme is activated by HBV infection. HBV has long been described as a “stealth” virus that elicits a limited hepatic IFN/ISG response, a feature that likely contributes to its ability to persist in hepatocytes^51-53^. Early-stage host responses to HBV infection are still debated, with some studies reporting minimal IFN/ISG induction during the initial phases of infection.

**Figure 4:**
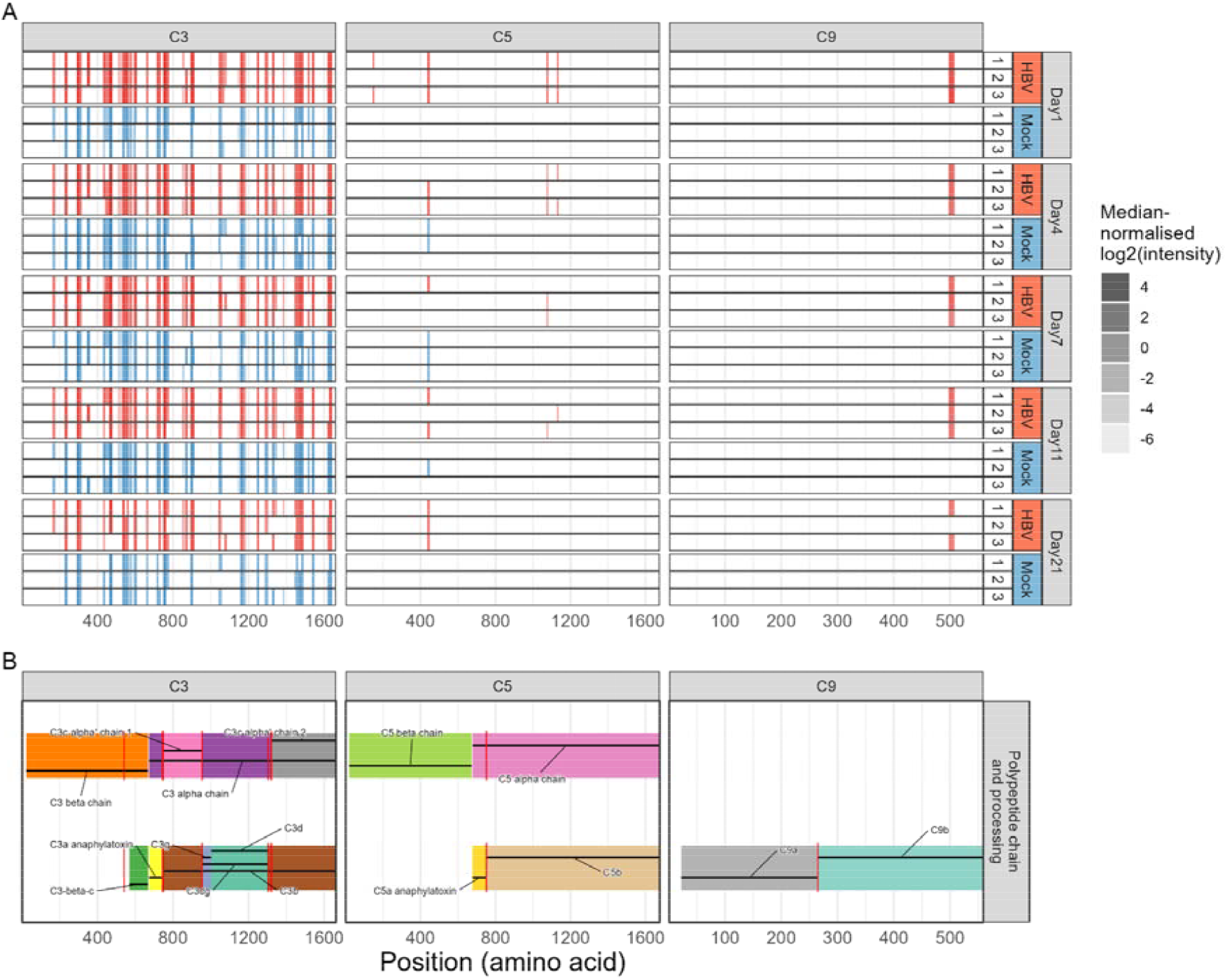
Altered complement profiles in HBV-infected HepG2-NTCP cells. Quantitative proteomics analysis and peptide mapping reveal time-dependent alterations in the levels of C5 and C9, but not C3, complement proteins. (A) Median-normalised log_2_(intensity) of peptides are shown for proteins C3 (left), C5 (middle), and C9 (right). Each replicate and time point is shown in ascending order for HBV- and mock-infected samples. (B) Reference polypeptide chains and processing. Complement chains (top) and fragments (bottom) are shown with corresponding labels. Complement protein cleavage sites are depicted as red vertical lines.

**Figure 5:**
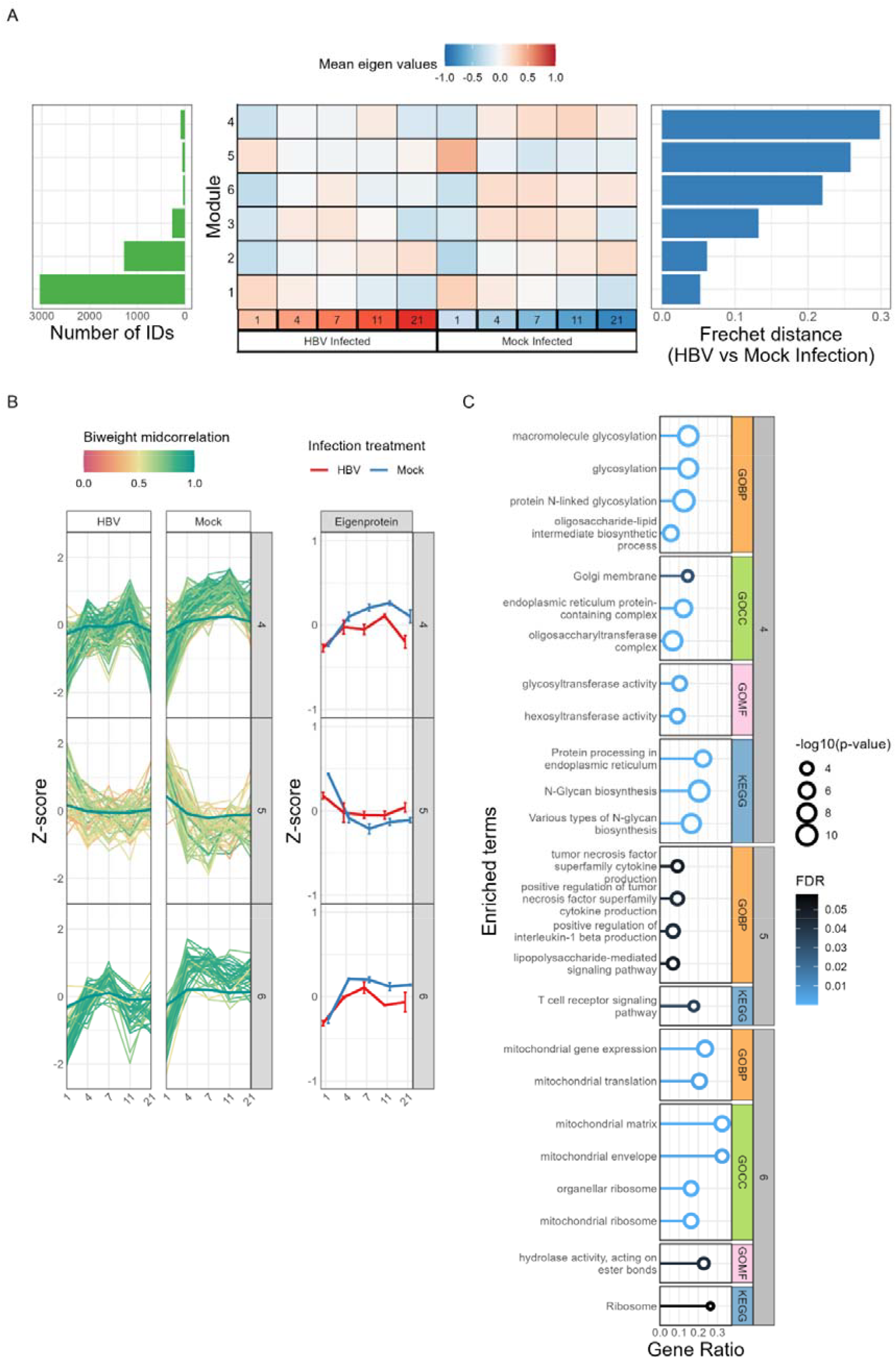
Weighted correlation network analysis identifies protein modules with differential abundance profiles in HBV infection. (A) Weighted correlation network analysis of log_2_-transformed and median-adjusted data containing imputed values. Each protein was Z-scored to standardise the protein abundances relative to their mean across all samples. Heatmap of module eigen proteins (middle), module membership (left) and Fréchet distances between mock and HBV-infected profiles (right) are shown. (B) Modules whose eigen protein expression profiles demonstrated higher Fréchet distances between mock and HBV-infected protein abundance profiles were subjected to further analyses (modules 4, 5, and 6). See **Fig. S7** for all clusters. Replicate data and module eigen protein values were mean averaged for each time point and treatment. Abundance profiles for each protein have been separated based on infection treatment (HBV and mock), and coloured based on biweight midcorrelation with the module eigen protein (left). Eigen protein profiles are also depicted alone (right). Error bars represent standard deviation of module eigen values for each timepoint and treatment. (C) Functional enrichment analysis of identified modules. Enriched gene sets for the GO terms and KEGG pathways, for each module, are shown, using the cellular proteome as background.

To assess whether HBV modulates this antiviral program at the protein level, we interrogated our global mass spectrometry–based proteomic dataset for ISG expression and regulation. We compiled a curated list of 573 ISGs implicated in antiviral defence and cross-referenced it with our proteomic data^54^. While only a small fraction of ISGs were significantly altered following infection, those that reached significance exhibited only modest fold-changes. These findings are consistent with the limited ISG induction previously reported for HBV and the notion of viral-mediated immune evasion^55-57^ **(Fig. S4)**.

WGCNA identified protein abundance changes in components of the secretory pathway and (protein) glycosylation upon HBV infection. As multiple steps in the viral replicative life cycle will involve ER–Golgi trafficking for protein processing and virion release, these findings are consistent with altered host secretory pathways. We speculate that reduced abundance of ER–Golgi-associated proteins may dampen cytokine secretion and thereby limit immune surveillance; however, this hypothesis requires experimental validation.

Studies involving human biopsies found common DAPs with *in vitro* studies such as ALDH1A1, ALDH2, ANXA4, APOA1 and GAPDH, underscoring the continued value of *in vitro* systems for studying proteome-wide alterations. In parallel, comprehensive efforts have catalogued HBV–host protein–protein interactions, generating increasingly detailed interactome resources^58^. However, these interaction maps largely represent static compilations derived from diverse experimental systems and do not capture the dynamic remodelling of the host proteome during authentic infection.

Our analyses were performed in an NTCP-expressing HepG2 hepatoma cell line, which does not fully recapitulate the complexity of primary hepatocytes or the multicellular liver environment *in vivo*. However, the controlled *in vitro* system enabled synchronised infection and systematic time-resolved sampling, an approach that is currently not feasible *in vivo* at comparable temporal resolution and proteomic depth. The limited induction of ISGs observed in our dataset suggests that, if present, interferon-driven responses do not result in substantial accumulation of antiviral effectors at the protein level. Our findings complement transcriptomic studies and provide an additional layer of insight into the regulation of early host responses during HBV infection. Taken together, the temporal resolution and proteomic depth of this study offer a framework for understanding hepatocyte behaviour during the initial stages of infection and support future mechanistic and *in vivo* investigations.

## ACKNOWLEDGMENTS

MA is funded by a Wellcome Four-year PhD Studentship in Basic Science 226811/Z/22/Z. The McKeating laboratory is funded by a Wellcome Investigator Award 200838/Z/16/Z, Wellcome Discovery Award 225198/Z/22/Z, Chinese Academy of Medical Sciences Innovation Fund for Medical Science, China (grant number: 2024-I2M-2-001-1). MB is funded by a King’s College Hospital Fund and the Chinese Academy of Medical Sciences Oxford Institute, CAMS Innovation Fund for Medical Sciences, funding code: 2024-I2M-2-001-1. ST, AD and BMK are funded by the CAMS Innovation Fund for Medical Sciences, funding code: 2024-I2M-2-001-1. This work was supported by the National Research Foundation of Korea (NRF) grant funded by the Korean government (MSIT) (No. RS-2024-00398073, NRF-2019R1A2C1090515).

## AUTHOR CONTRIBUTIONS

AK, EJ and MW developed the cell line and HBV infection model. SH and AD processed the samples and prepared them for mass spectrometry analysis. SH and IV performed the mass spectrometry experiments. AD and MA analysed the proteomics data. MA, MB and ST contributed to data interpretation. MA, MB and AD drafted the manuscript. All authors reviewed and edited the manuscript. BMK, JAM and AD contributed to the conceptualisation and overall design of the study.

## Data Availability Statement

The mass spectrometry proteomics data generated in this study have been deposited to the ProteomeXchange Consortium via the PRIDE partner repository under accession number PXD074747. All other data supporting the findings of this study are available within the article and its Supporting Information.

## Ethics Statement

This research did not involve human or animal participants.

## Supporting Information

The following supporting information is available free of charge at ACS website http://pubs.acs.org

**Figure S1:**
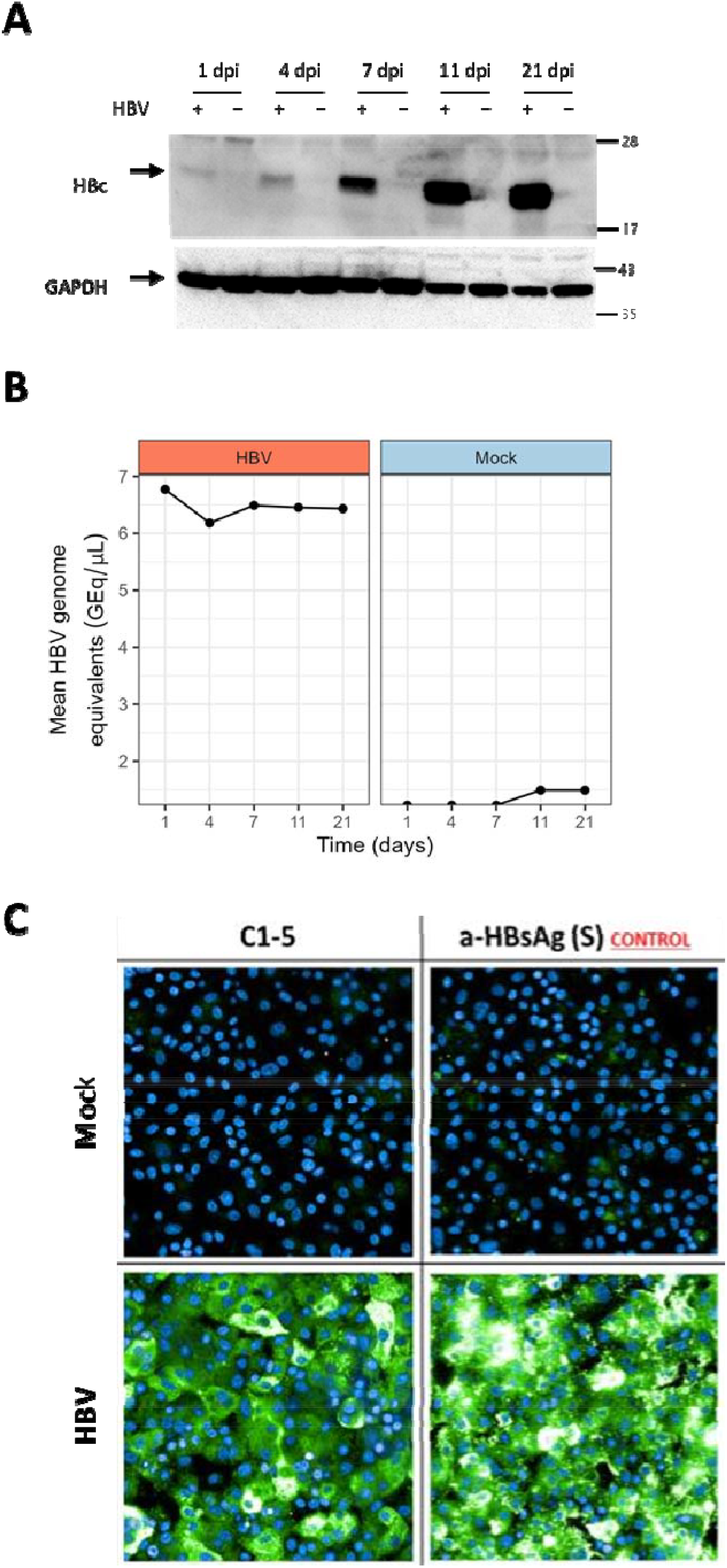
Validation of HBV infection. (A) Western blot validation of samples subjected to proteomic analysis. HBV core protein (HBc) and GAPDH loading control are shown for infected and mock samples. (B) PCR-based quantification of HBV genome equivalents for infected and mock-infected samples at each time point. (C) Immunofluorescence microscopy images of HBc and HBsAg expression at 21 dpi.

**Figure S2:**
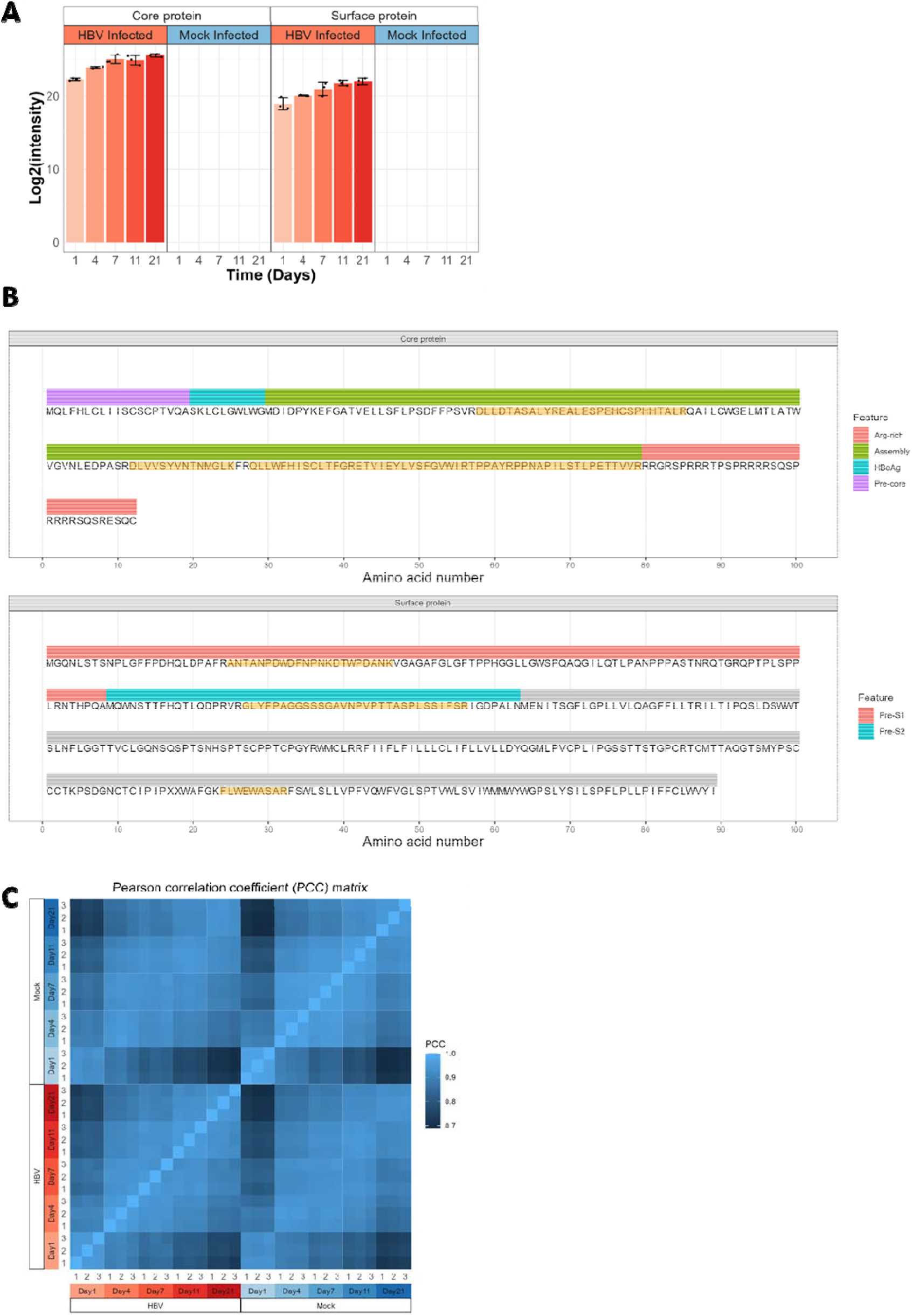
Proteomic analysis of the HBV infection model. (A) Mean log_2_-intensity values for Core (C, left) and Surface (S, right) proteins from HBV in each condition (time and treatment groups). Error bars represent standard deviations. (B) Mapping of HBV-derived tryptic peptides. Identified peptides mapping to the shared precore/core coding sequence (C) and the HBV surface (S) proteins. (C) Pearson correlation coefficients between biological replicates, time points, and treatment groups of log_2_-transformed, median-normalised, and imputed data.

**Figure S3:**
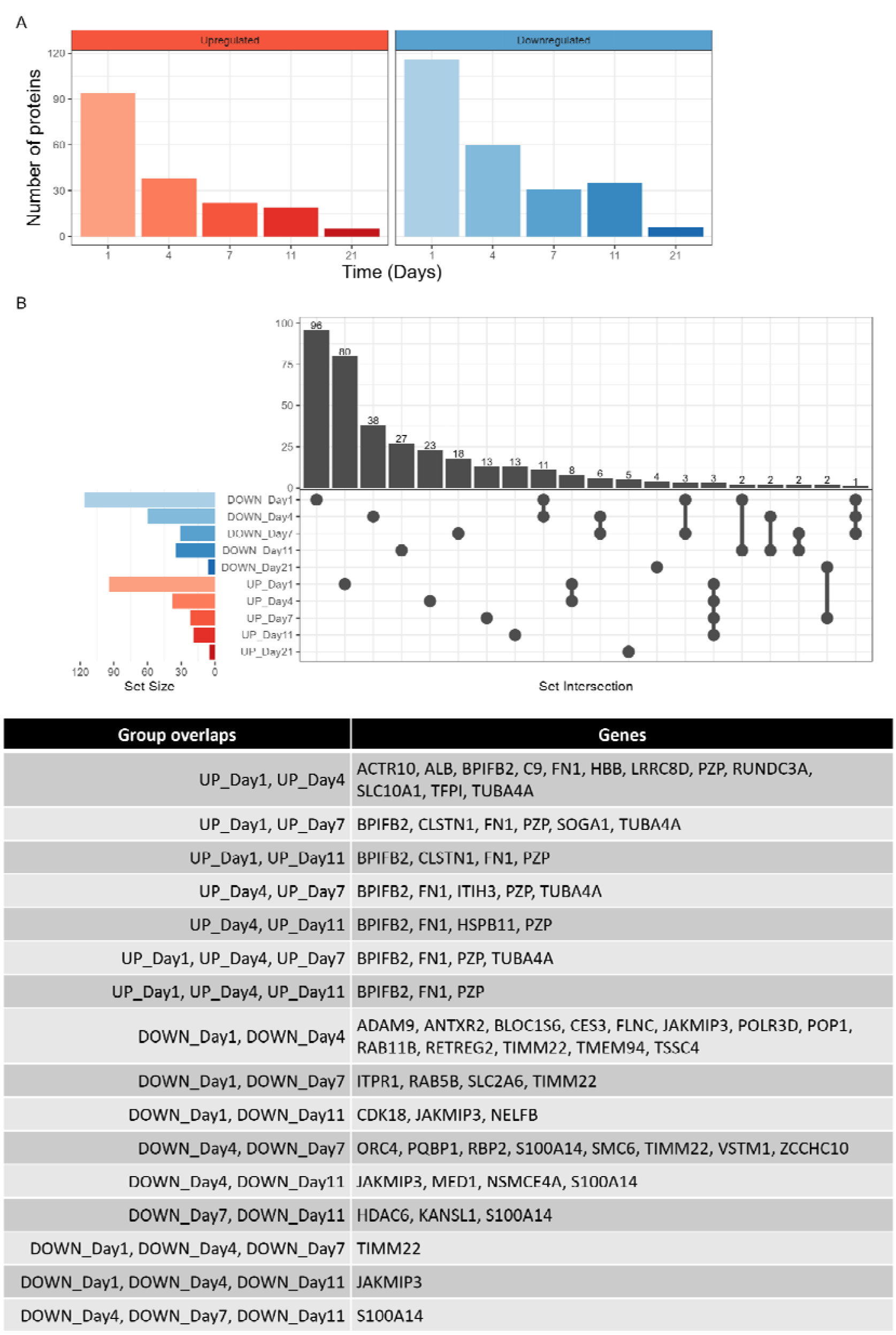
Summary of DAPs identified using moderated t-test. (A) Total number of DAPs identified for each time point. (B) Upset plot showing the number of overlapping proteins identified as differentially abundant. Overlapping groups and the corresponding DAPs are listed in the table below.

**Figure S4:**
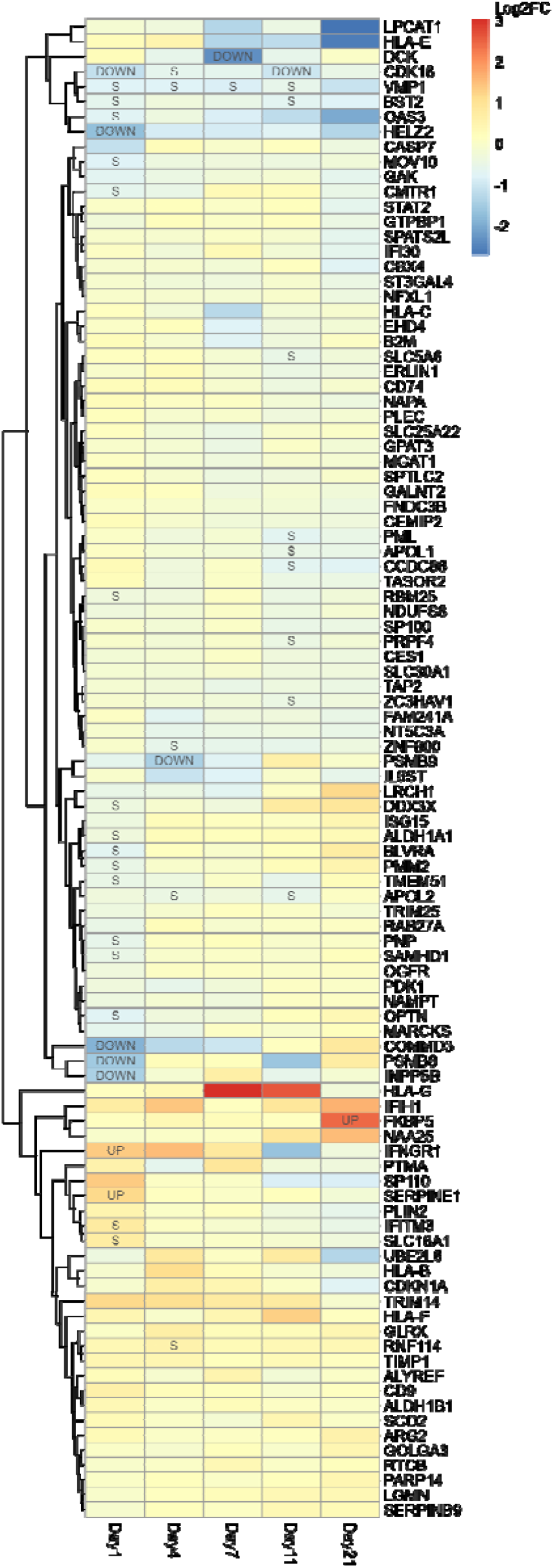
ISG protein abundance changes during HBV infection. Heatmap of the top 100 interferon-stimulated genes based on the highest |log_2_FC| across all time points. A dendrogram represents hierarchical clustering of protein log_2_FC based on Euclidean distance. UP = FDR < 0.1, log_2_FC > 1; DOWN = FDR < 0.1, log_2_FC < -1; S = FDR < 0.1, |log_2_FC| < 1.

**Figure S5:**
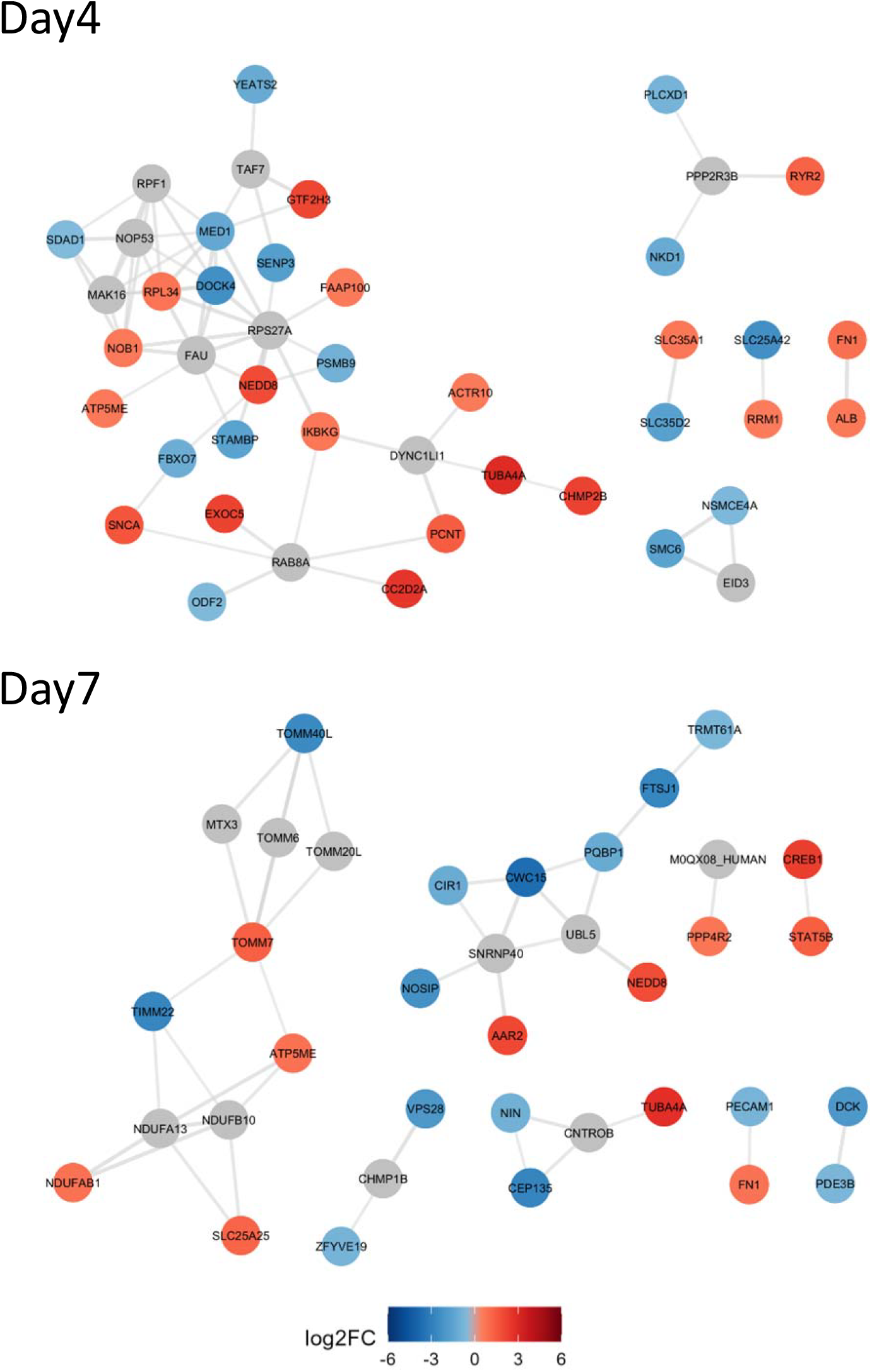
Network analysis of significantly altered proteins during HBV infection. Protein–protein interaction networks of significantly upregulated and downregulated proteins (FDR < 0.1, |log_2_FC| > 1) at 4 dpi (A) and 7 dpi (B). Node colour represents the log_2_FC in protein abundance relative to mock controls, as derived from quantitative proteomics. Grey nodes represent additional interactors (not identified as DAPs), as determined by STRING. Edges denote high-confidence functional and/or physical associations (minimum required interaction score > 0.7). Interaction data were retrieved from the STRING database (v12.0) using the stringApp in Cytoscape (v3.10.0). Isolated (unconnected) nodes were removed for clarity.

**Figure S6:**
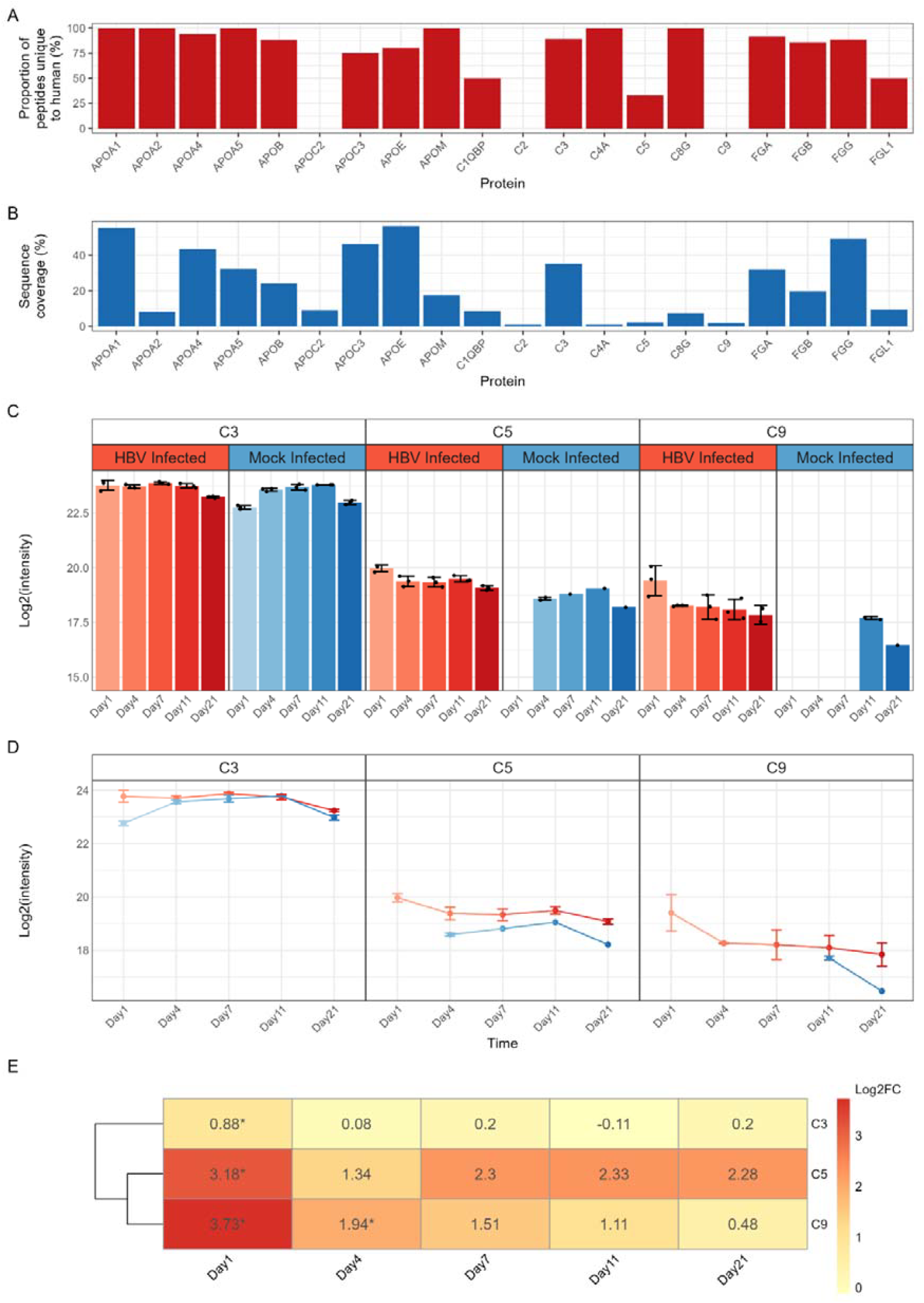
Peptide uniqueness, sequence coverage, protein-level trajectories, and statistical significance for proteins of interest. (A) Human sequences for complement-, fibrinogen-, and apolipoprotein-associated proteins were aligned to bovine homologs, and identified peptides were mapped to bovine counterparts. The proportion of peptides unique to human sequences for each protein were calculated. (B) Sequence coverage of proteins as percentage of amino acids detected in peptides. (C) Mean Log2(intensity) values for C3, C5, and C9. Error bars represent standard deviation. (D) Protein-level trajectories of C3, C5, and C9. Error bars represent standard deviation. (E) Heatmap of moderated t-test values of C3, C5, and C9, comparing HBV to mock infection treatment. Numbers indicate Log2FC, and asterisks indicate *FDR < 0.1.

**Figure S7:**
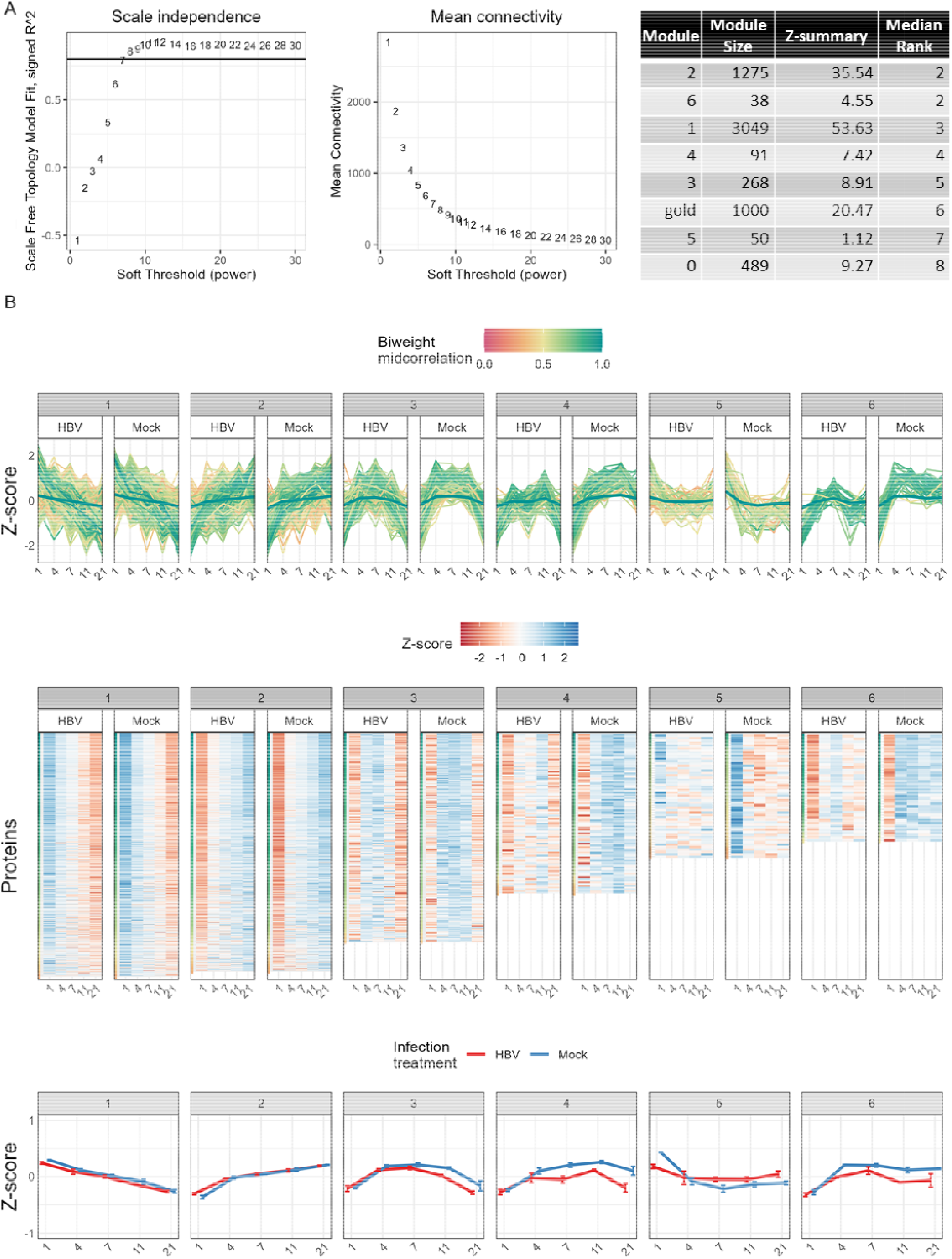
Protein abundance correlation profiles during HBV infection. Weighted correlation network analysis of log_2_-transformed and median-adjusted data containing imputed values. Values were Z-scored to standardise the abundance levels relative to their collective mean across all samples. (A) Soft-thresholding optimisation identified a power of 12 as an appropriate parameter, based on scale independence and mean connectivity. Module preservation statistics are shown on the right, where the ‘gold’ module represents a random, control module. (B) All modules (clusters) are shown. The top panels show eigen protein and individual protein Z-score profiles for mock and HBV-infected samples, the middle panels are heatmap representations, and the bottom panels indicate eigen protein abundance profiles for each treatment over time. Error bars represent standard deviation of eigen values per time point and treatment within the module.

